# Sex differences in parental response to offspring begging are associated with pair bond strength across birds

**DOI:** 10.1101/2021.10.14.464423

**Authors:** Shana M. Caro, Karleen Wu, Hans A Hofmann

## Abstract

Mothers, fathers, and offspring regularly clash over how much care offspring receive. Offspring beg to solicit for more resources—but how much begging is rewarded can depend on who is listening. While both parents benefit from provisioning offspring, each would benefit from their partner shouldering more of the burden of care, leading to sexual conflict. Additionally, if the costs and benefits of provisioning differ by sex, parent-offspring conflict should vary by sex. How these evolutionary conflicts influence sex differences in parent-offspring communication is unknown. To determine whether the sexes differ in their response to offspring signals, we conducted a meta-analysis on 30 bird species, comparing responsiveness to social and physiological traits affecting conflict. We found that a species’ typical pair bond strength predicts whether males or females respond more to offspring begging. In species with stable and/or monogamous bonds, and thus lower sexual and paternal-offspring conflict, males’ provisioning effort is more strongly correlated with offspring begging than females’. The opposite holds for species with weak pair bonds: females respond more to begging, perhaps compensating for males’ lower responsiveness. These results demonstrate that sex differences in parental care can arise via sex differences in parent-offspring communication, driven by evolutionary conflicts.

## Introduction

Parental care is a wellspring of evolutionary conflicts. Caring for young can be costly, and each parent would have higher fitness if their mate did more of the work—leading to sexual conflict over parental care (1–8). For example, in species such as the Kentish plover (*Charadrius alexandrines*), offspring initially require the care of two adults to survive, but eventually one adult becomes sufficient (9). Whichever parent abandons the nest first can mate again with a new partner, potentially doubling their reproductive output for the year, while the remaining parent must continue caring for the original brood (9). Many factors affecting the degree of sexual conflict over care and how much care each sex provides have been identified by theoretical models, notably the strength of pair bonds (2,10–13), sex ratio (3,14,15), and sex differences in the cost or ability to provide care (6,16,17).

However, such models frequently do not explicitly consider another player that impacts how much care each sex provides: the offspring themselves. Offspring and parents are frequently at odds regarding the optimal level of care. Offspring may benefit from maximizing the care they receive, but parents benefit by maximizing their lifetime reproductive success (18,19). Offspring across many taxa have evolved to entice their parents to increase their provisioning effort via “begging” signals, such as vocalizations, postures, and brightly colored mouths (20). How offspring signal and how parents respond to those signals depends on the degree of parent-offspring conflict over parental investment, which in turn is impacted by factors such as promiscuity, pair bond stability across reproductive bouts, the number of offspring produced per brood and over the parent’s lifetime, and how relatively abundant resources are (19,21–23).

These two axes of familial evolutionary conflicts—parent-parent and parent-offspring—intersect in how much each sex increases their parental effort in response to offspring signals (Figure 1). In nature, the extent and direction of a sex difference in how parents respond to begging varies from female-biased (e.g. 24–26), to no-bias (e.g. 27,28), to male-biased (e.g. 29,30). Why this natural variation exists is unclear. One possibility is that there is no true sex difference in responsiveness to begging, and studies showing otherwise have captured rare exceptions to the rule or are statistical anomalies. Another possibility is that males and females differ in some species but not others, and that this pattern can be explained by the same factors that influence other aspects of conflict over parental care. Parents’ response to offspring signals is thus an excellent lens through which to investigate evolutionary conflicts in species with biparental care.

**Figure 1.**
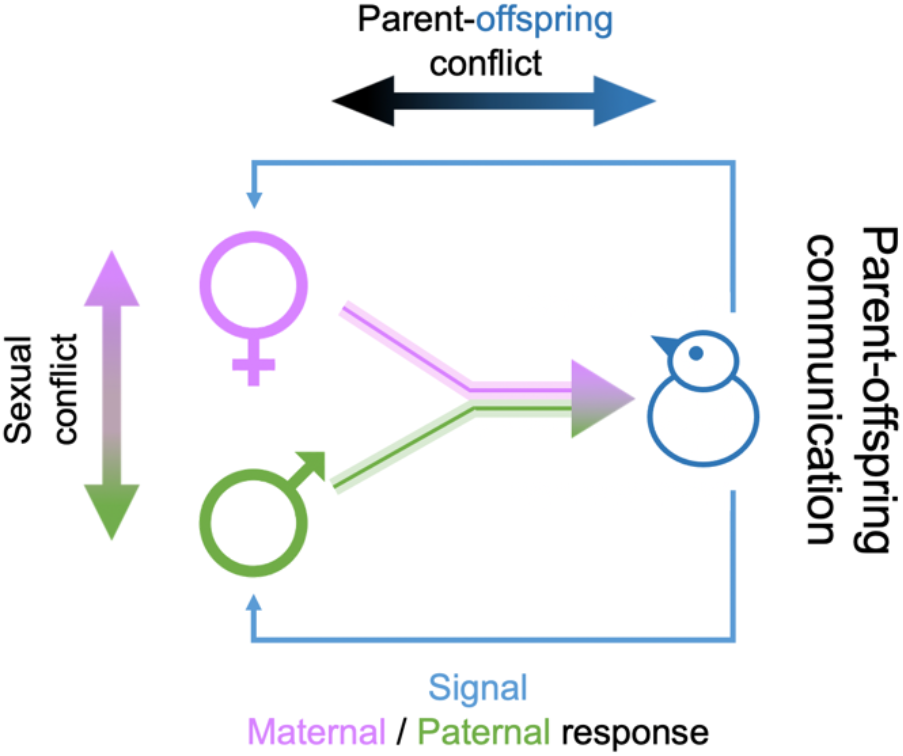
How much care parents provide is a battleground for both parent-offspring and sexual conflict, partly mediated by parent-offspring communication. Males (green) and females (purple) would benefit from having their mate provide more of the care necessary for rearing offspring, while offspring (blue) would benefit from receiving more care than parents’ optimal level. Offspring signal for more resources (blue arrows), and parents can respond (or not) by increasing their parental effort. Solid lines indicate each parent’s desired investment, and transparent lines indicate their potential investment—and how much their mate and offspring might prefer them to invest.

To investigate whether—and why—males and females differ in their provisioning response to offspring begging, we conducted a comparative meta-analysis across 30 species of birds. Comparative approaches are invaluable for explaining broad evolutionary patterns, because within-species variation in the relevant traits may not be sufficient to generate differences in parent-offspring communication, while the greater variation across species is sufficient. We considered responsiveness to behavioral begging signals as the change in total provisioning effort: e.g., how many feeding trips does each sex make when offspring beg louder or how much more food is provided when offspring spend more time begging. While parental care can take many forms—cleaning, housing, defending, thermoregulating—here, we focus on increased food provisioning because this response to begging is the most well-studied (20). We restricted our study to birds because at least 80% of birds show biparental care (31) and begging is most well-studied in birds (20). First, we assessed whether a sex difference exists, and hypothesized that females increase their provisioning efforts more in response to begging than males, since females are typically the more caring sex (13). We next investigated whether sex differences co-vary with factors known to influence sexual and/or parent-offspring conflicts: i) pair bond strength (a categorical factor combining promiscuity and the duration of pair bonds); ii) divorce rate; iii) extra-pair paternity (promiscuity) rate; iv) strength of sexual selection (sexual dimorphism in size and plumage); v) baseline corticosterone hormone (stress) levels while caring for nestlings; and vi) ecological conditions. We hypothesized that species with weak (unstable, promiscuous) bonds would have more conflict between the sexes due to decoupled future fitness and changes in average male relatedness to the brood, and that males would experience more parent-offspring conflict because of their reduced paternity certainty, leading to lower male responsiveness (16,22,32). Conversely, species with strong (stable, monogamous) bonds would have more alignment of male and female future fitness, and fathers would have more paternity certainty, leading to higher male responsiveness and lower sex differences (2,12,23,32). Alternatively, because increased male care may allow females to breed again more quickly (as seen across mammals, 33), we hypothesized that males would be more responsive than females mostly in species with strong bonds. Sexual dimorphism in size and plumage is associated with higher levels of sexual conflict (2,10,34). We hypothesized that more physically dimorphic species would show more dimorphism in responsiveness. The glucocorticoid hormone corticosterone is a well-known indicator of physiological stress, and may thus be indicative of the costs of care (6,17,35). We hypothesized that the costlier care is for a sex, as shown by stress hormone levels, the less that sex would respond to begging. Finally, ecological conditions that influence the relative abundance of food are associated with changes in parent-offspring communication regarding within-brood food allocation, and we investigated their effects (21).

## Methods

### Data collection

We conducted a literature search on Google Scholar using the keywords “beg”, “begging”, “parent-offspring”, “communication”, and “bird” (see Supplementary Figure 1 for PRISMA flowchart detailing data collection). We performed forward citation searches on all studies. We included all papers with any measure relating provisioning effort to behavioral begging, where the measure was separated by sex. If males and females were pooled in reported analyses but it seemed likely the authors had recorded sex, we emailed authors to request additional data. We included six unpublished datasets, graciously provided by the following researchers: Drs. Alejandro Cantarero, Niels Dingemanse, Arne Iserbyt, Kirsty MacLeod, Mark Mainwaring, Ariane Mutzel, and Jonathan Wright. This resulted in a data set of 156 effect sizes in 48 studies on 30 species (Supplemental Table 1). For 28 species, we found data on both males and females, but two species only had data on one sex. A list of excluded studies and rationale can be found in Supplemental Table 2.

### Estimating effect sizes

We measured responsiveness to begging as the effect of behavioral begging (e.g. vocalization rate or postural intensity) on total provisioning effort (e.g. feeding rate or amount of food provided). We found no impact of data type on effect sizes (see Confounding Factors section). We only included effect sizes for the relationship between begging and total provisioning amount, rather than food allocation within a brood, as these represent fundamentally different aspects of parental provisioning. We transformed test statistics, or raw means and variances from data or figures if no test statistics were provided, into standardized effect sizes (Fisher’s Z-transformed correlation coefficient), which are normally distributed, independent of the scale of the original measurement, and easy to interpret (36). Positive correlation coefficients indicate that the more offspring beg, the more parents feed. Negative correlation coefficients indicate that the more offspring beg, the less parents feed. Variance of the Z-transformed correlation coefficient is ^1^/(n-3), where n is the number of broods in the original test statistic. We assumed that all measures of begging and feeding were capturing aspects of the same biological phenomenon, and therefore included all test statistics in our analyses.

We quantified the within-species sex difference as the difference in responsiveness to begging for males and females of the same species: the mean male Z-transformed correlation coefficient minus the mean female Z-transformed correlation coefficient. Thus, negative values indicate that females respond more than males in that species, while positive values indicate that males respond more than females in that species. We used this measure rather than including a random slope for sex within species, because random slopes require multiple observations per level of a random effect, and some species-sex combinations had only one effect size measure. For cooperatively breeding species (n = 7), we considered the responsiveness of only the presumed breeding male and female.

### Data on explanatory factors

To explain any sex differences we might find, we collected data on a variety of species’ traits, and looked for (i) interactions between sex and that trait on Z-transformed correlation coefficients; and (ii) an impact of that trait on the within-species sex difference. For mating system, we initially recorded the species’ promiscuity (the mean rate of extra-pair paternity both by brood and by offspring) and pair bond stability (divorce rate: the likelihood of not mating again with the same individual in the next year). Because exact values for these traits were not available for all species, we employed a categorical measure of social bond strength developed by Tobias et al 2016 (37), which combines multiple aspects of social bond strength, including mate fidelity and divorce rates. Specifically, species with **weak** pair bonds (n = 11 species in the present study) are characterized by low mate fidelity and high divorce rates (>50% per annum), whereas species with **strong** bonds (n = 19 species) display high mate fidelity and low divorce rates (<50% per annum).

We investigated whether three aspects of bird physiology correlated with sex differences in responsiveness: plumage dimorphism following (10), a proxy for sexual selection; skeletal size dimorphism (via tarsus bones), a putative measure of either sex differences in the cost of care or sexual selection; and baseline corticosterone levels during young care, a putative measure of sex differences in the cost of caring for young (38).

Because environmental predictability and quality have been shown to influence parent-offspring communication, we recorded whether effect sizes were from times when environmental conditions were worse than normal, normal, or better than normal; and whether species are from predictable environments (clutch-adjusting) or unpredictable (brood-reducing) environments (21).

### Statistical analysis

We analyzed how males and females respond to begging and the degree and direction of the within-species sex difference using Bayesian linear mixed models with Markov chain Monte Carlo methods, using the MCMCglmm package in R (39,40). We weighted models by sample size, and controlled for phylogeny (all models) and for repeated measures from the same study and species (across-species models) (36,41). Sample size was the number of broods used in the original test statistic. We used informative priors, given previous work showing moderate phylogenetic signal on this communication system, and given the conservative assumption that males and females from the same species would be more similar to each other than to other species. Models with uninformative priors yield similar results (e.g. results change from post.mean = −0.17, pMCMC = 0.036 to post.mean = −0.16, pMCMC = 0.048). Because we had reduced sample size for many of our explanatory factors, we generated separate models for each factor. We find qualitatively similar results when we include all factors in the same model, though our sample size drops to 11 species (results not shown). Models had 3,000,000 iterations, with a burn-in of 1,000,000 and were thinned every 1,000. Model convergence was assessed visually by looking at variance-covariance and solution plots. We generated models on 20 random phylogenetic trees downloaded from BirdTree.org, using Erikson and Hackett backbones (41). Reported results are the average across these models.

We analyzed the heterogeneity of our data and assessed publication bias using the metafor package in R (42). We used Egger’s test for funnel plot asymmetry to investigate whether published data is biased by the “file drawer” problem of not publishing null results (Supplemental Figures 2 and 3).

Example R code for each of the main models presented here can be found in the supplemental material, and the full code and dataset for all models can be found at github.com/shanacaro/SexDifferencesParentalResponseBegging.

### Confounding factors

To ensure that our findings were not driven by confounding effects of the methodology used in the original studies, for each effect size we recorded data on experiment type (observational or experimental); whether the effect size was estimated from figures or generated from test statistics or data (estimated or not); begging mode (audio, postural, or a combination); begging variable type (continuous or dichotomous); feeding variable type (continuous or dichotomous); whether chicks were food deprived (yes or no); whether chicks or parents were food supplemented (yes or no); whether brood size was manipulated (yes or no); and whether playbacks of begging calls were used (yes or no). None of these methodological factors had any influence on the Z-transformed correlation coefficient between begging and feeding (MCMCglmm phylogenetically controlled and weighted regression: pMCMC > 0.20; Supplementary Table 3).

### Heterogeneity

We measured the heterogeneity in our effect sizes with I^2^, the proportion of observed variance due to true differences in effect sizes rather than measurement errors (36,43,44). I^2^ was calculated by dividing the summed variance attributed to phylogeny, study, species, and unit, by the total variance in the data (variance attributed to measurement error, phylogeny, study, species and unit). The higher the I^2^ value, the more indication that observed variance is due to true differences, with 25%, 50% and 75% as low, moderate, and high benchmarks. We also report Cochran’s Q statistics, which tests the probability that variation is greater across studies compared to within studies (36). Larger Q values indicate that studies do not share the same true effect size.

### Heritability

We measured heritability as the variance that can be attributed to the phylogeny in a null model with no fixed effects, with phylogeny, study and species as random effects, and weighted by sample size. We also measured heritability in each of the models we generated with fixed effects. This was calculated by dividing the variance attributed to phylogeny by the overall variance in the data. The phylogenetic signal in the null model was 14%.

## Results

First, we asked whether across species, males and females vary in how they respond to offspring begging, but we found that they are equally responsive. There is a significant correlation between begging and total provisioning effort for both sexes (female Z-transformed correlation coefficient = 0.43, 95% CI 0.21 to 0.67, pMCMC < 0.002; male Z-transformed correlation coefficient = 0.46, 95% CI 0.23 to 0.68, pMCMC < 0.002; sex difference = 0.02, 95%CI −0.08 to 0.09, pMCMC = 0.71; Figure 2). Phylogeny accounts for 16% of the variance in how males and females respond.

**Figure 2.**
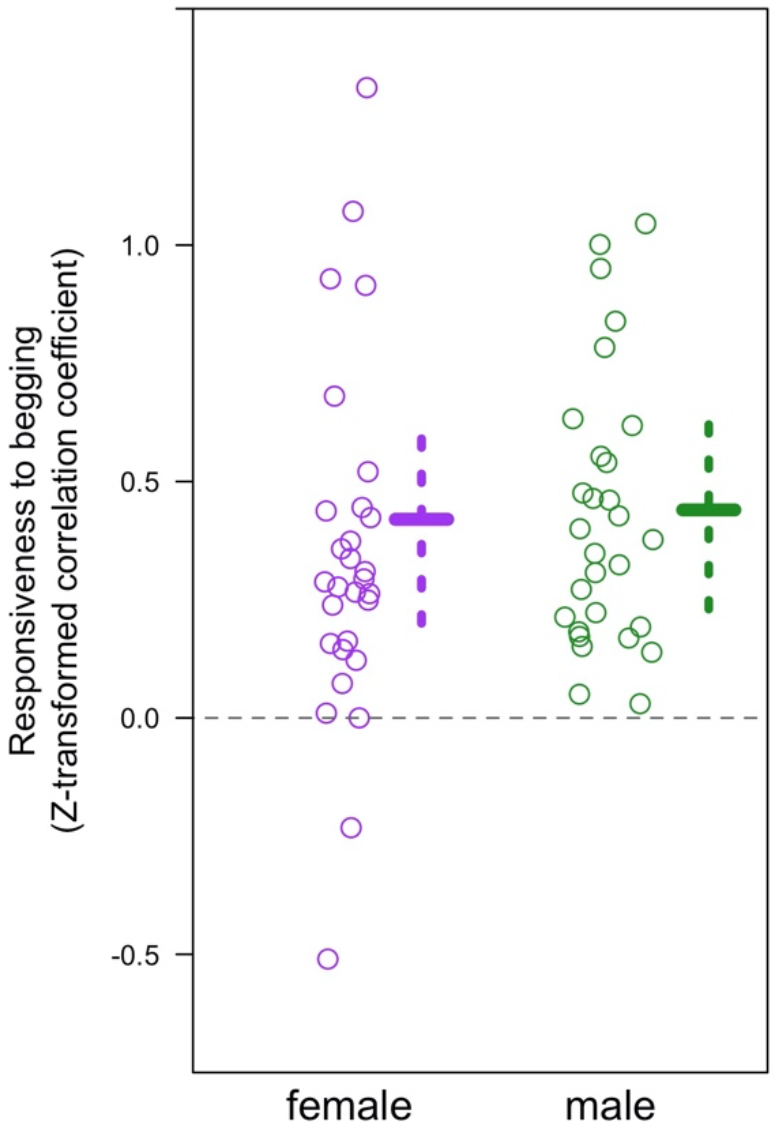
Across species, males and females do not differ in their responsiveness to offspring begging (pMCMC = 0.71). Data points represent the mean Z-transformed correlation coefficient for a species. Solid and dashed lines represent the results from the MCMCglmm models: post.mean and 95% credible intervals (purple = female, green = male). N = 30 species.

Although the overall pattern shows no difference between the sexes in their responsiveness to offspring, there is considerable variance concealed within a species. As predicted, species differ in whether males or females respond more to offspring begging. In roughly a third of species, the female responds most strongly to begging, in another third there is little sex difference, and in the last third the male responds most strongly (Figure 3). We found high heterogeneity in the mean effect size per sex in each species (I^2^ = 76.6%, Q_57_ = 197.8, p < 0.0001), indicating that males and females in different species respond differently to begging. We found no evidence of publication bias (Egger’s test z = 1.28, p = 0.20).

**Figure 3.**
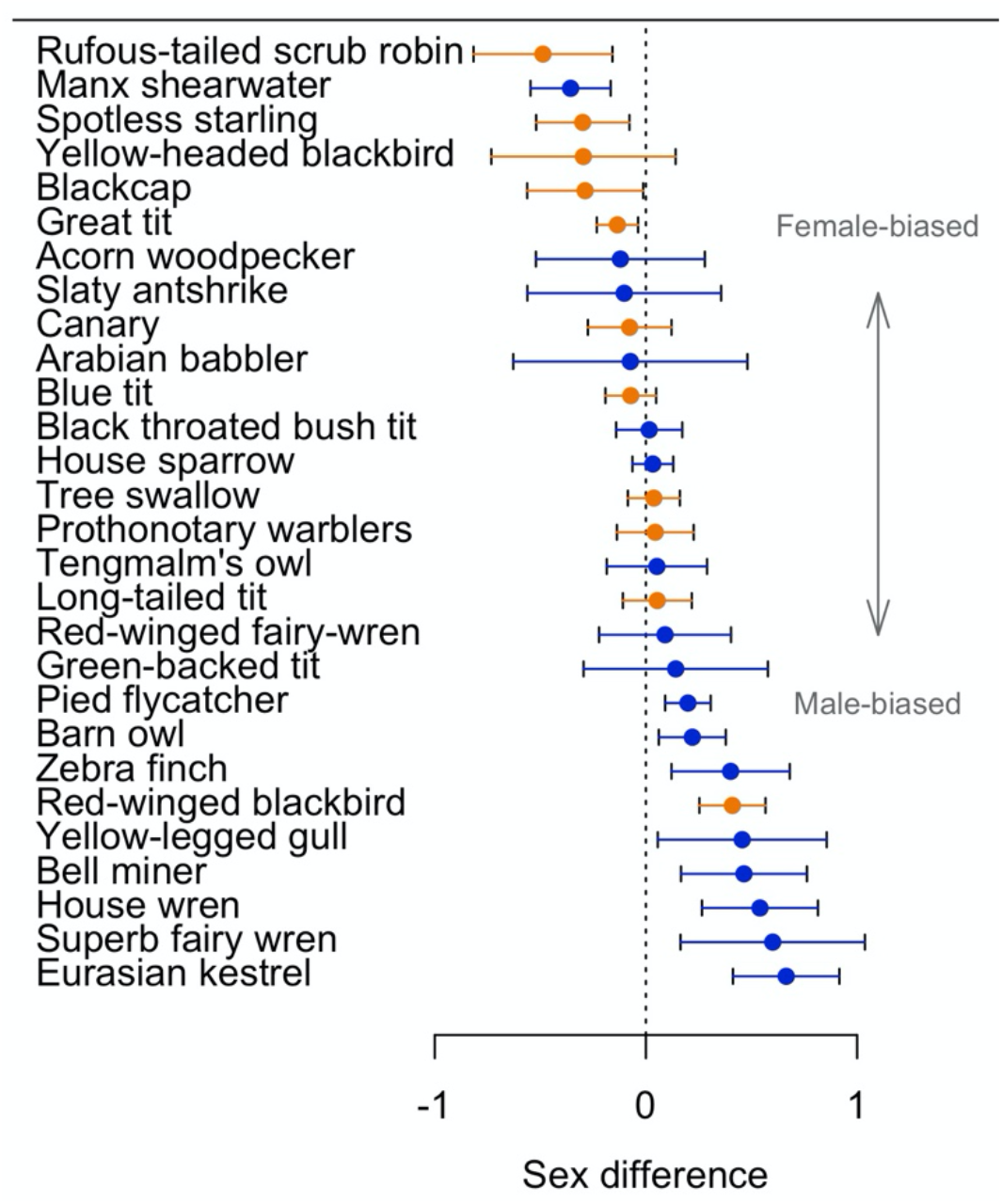
Species vary in the degree and direction of the sex difference in how parents respond to offspring begging. The forest plot shows the means and 95% confidence intervals for the within-species sex difference (male mean Z-transformed correlation coefficient – female mean Z-transformed correlation coefficient). Negative values indicate females respond more, and positive values indicate males respond more. For example, male pied flycatchers have a mean Z-transformed correlation of 0.27 and females have a mean Z-transformed correlation of 0.07, so the sex difference for this species is 0.20. Data points are colored by pair bond type: weak pair bonds are represented by orange, and strong pair bonds by blue. N = 28 species.

Given this heterogeneity across species, we tested the hypothesis that how each sex responds to begging is associated with variation in the strength and stability of pair bonds. Indeed, we found a significant interaction between sex and social bond type (post.mean = −0.16, 95% CI −0.33 to - 0.002, pMCMC = 0.048*; Supplemental Table 4), suggesting that females are more responsive to begging in species with weak social bonds than in species with strong social bonds, while the opposite pattern holds for males (Supplemental Figure 4).

**Figure 4.**
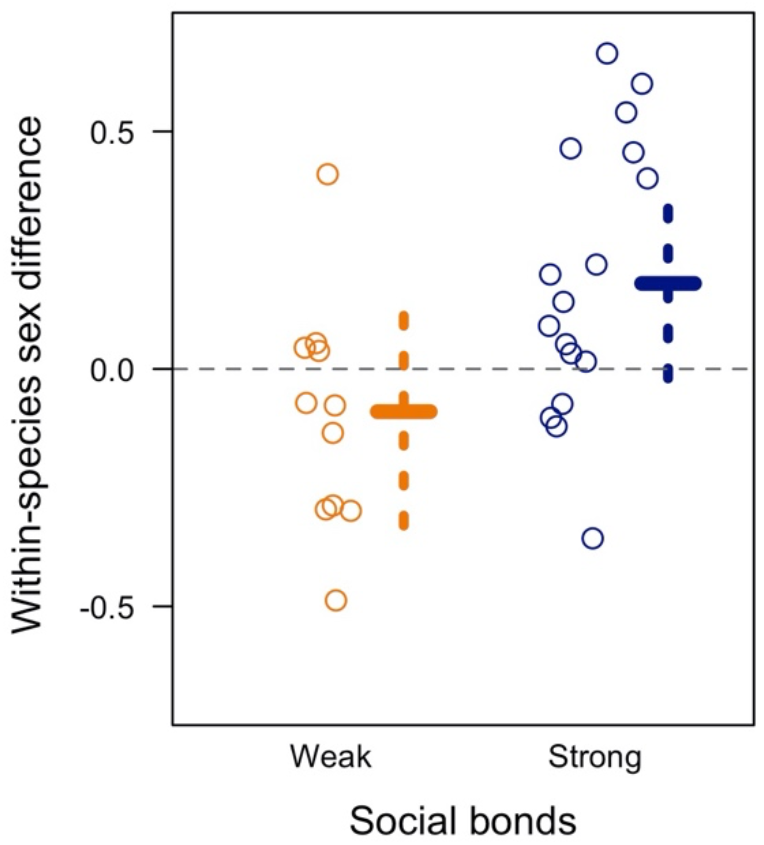
Within-species, pair bond strength determines which sex responds more to begging. Within-species, males are more responsive to begging in species with strong social bonds, while females are more responsive to begging in species with weak social bonds (pMCMC = 0.047*). Data points represent the sex difference in parental responsiveness. Negative values indicate females are more responsive, positive values indicate males are more responsive. Solid and dashed lines represent the results from the MCMCglmm models: post.mean and 95% credible intervals. N = 28 species (11 with weak social bonds, 17 with strong social bonds).

Next, we asked whether a species’ typical mating system determines the degree and direction of the within-species sex difference in responsiveness. We found that within species with strong pair bonds, males are more responsive to begging than females, whereas within species with weaker pair bonds, females are more responsive than males (estimate = 0.26, 95% CI = 0.004 to 0.53, pMCMC = 0.047*, Figure 4, Supplemental Table 5). This within-species pattern mirrors the pattern seen across species. Phylogeny accounts for 17% of the variance in the within-species sex difference. We found no evidence of publication bias for within-species sex differences in responsiveness (p = 0.54, Supplemental Figure 3).

No other explanatory factors significantly influenced how males and females respond to begging (Supplementary Table 6, all pMCMC > 0.10). We did not find an effect of tarsus dimorphism (24 species), promiscuity as a continuous measure (n = 21 species), divorce rate as a continuous measure (n = 16 species), plumage dimorphism (n = 12 species), environmental predictability (n = 30 species), or environmental quality (n = 30 species). Finally, we found that males in species with higher baseline levels of corticosterone during care for young showed a trend of decreasing responsiveness to begging (z = −2.4, p = 0.078), while females tended to be most responsive to begging when baseline corticosterone levels are at intermediate levels (corticosterone z = 2.97, p = 0.059; corticosterone^2^ z = −3.0, p = 0.057). However, neither effect was significant when controlling for phylogeny, likely because of a lack of statistical power: baseline corticosterone levels during care for young were available for only six species.

## Discussion

Our comparative study found that males and females differ in how they respond to offspring begging, although this sex difference is obscured when looking across species. Within species, whether it is males or females that increase provisioning effort more in response to begging depends on pair bond type. In species with strong and stable pair bonds, i.e. with lower scope for sexual conflict, males are more responsive than females. In species with weak and unstable pair bonds, females are more responsive than males. Pair bond strength is implicated in both sexual conflict—by linking or decoupling future reproductive success for males and females—and parent-offspring conflict—by varying expected relatedness between males and offspring. Our results therefore suggest that males and females may have evolved different levels of responsiveness to offspring signals based on variation in sexual conflict and parent-offspring conflict, due to differences in pair bond type at the species’ level.

Sexual conflicts over parental care arise when males and females would each have higher fitness if their mate did more of the work of rearing offspring (18,32). In species with biparental care, many adaptations have evolved in response to sexual conflict over parental care. There may be a sexual division of labor, where each sex specializes on particular tasks and/or offspring (27,32,45). Parents may negotiate care indirectly through monitoring offspring (46–48) or directly through coordinating feeding visits or vocal communication (12,49,50). Males may also use parental care as a form of sexual signaling, if females preferentially mate with good fathers, as is found across fishes (51). Our study provides evidence for an additional avenue through which sex difference in parental care emerge: parent-offspring communication. By varying how much they respond to offspring signals of need, parents can: i) create a sex difference in parental effort, if both sexes care equally in the absence of begging; ii) exaggerate a sex difference, if the more responsive sex also provides more care in the absence of begging; or iii) ameliorate a sex difference, if the more responsive sex provides less care in the absence of begging. Future experiments could investigate baseline sex differences in parental effort in the absence of begging, as in (52) which deafened ring dove parents to remove the effect of begging vocalizations. Future studies could also investigate whether within-species variation in pair bonds is sufficient to influence maternal and paternal provisioning patterns, or if this pattern only emerges at the species level (16). Additionally, offspring have been observed begging differently to their mother and father (53), though it is unclear how widespread this phenomenon is. Finally, parents care for offspring in a myriad of ways other than provisioning; for instance, parents may respond to begging by warming their chicks, and future studies could examine sex differences in begging and other aspects of parental care.

Our study has implications for the evolution of parent-offspring communication, and for the evolution of communication systems more broadly. Firstly, we found that males and females can vary in how they respond to offspring begging signals. Models of parent-offspring communication, however, typically only consider one parent. Parental care models have looked at interactions between parents, but these typically do not include offspring effects (32,48). Future models could therefore include three classes—offspring, mothers, and fathers—to investigate how begging and parental care is predicted to evolve when there are multiple responder classes that may or may not be correlated in how they respond to signals. Additionally, although females are always equally related to their offspring, we found that they are more responsive to offspring in species with typically weak pair bonds. Females in such species seem to be compensating for unresponsive fathers by increasing their own responsiveness to offspring. This finding implies that communication between two individuals, a parent and an offspring, can be impacted by a third individual, the other parent. While parent-offspring signaling models have explicitly taken the abiotic environment into account (for example 54), they have been less likely to consider the social environment. Our results indicate that the social environment matters, perhaps just as much as the abiotic environment. Explicit mathematical models on sexual conflict and parent-offspring communication, rather than verbal models, will be necessary to clarify these relationships (23).

Begging is not just seen in birds; it has also been observed in burying beetles, poison frogs, ants, mice, humans, and more (20). Studies across the animal kingdom have shown that social systems and the costs and benefits of care determine the level of parental care (18,33,51). I t is less well understood how maternal and paternal care varies *flexibly* based on signals from offspring in other taxa with biparental provisioning. For example, a recent study across three species of burying beetles found that how much each sex of each species provisioned their begging offspring when their mate was nearby varied, but for unknown reasons (24). Other factors may be at play in different taxa, such as dispersal patterns: in the cooperatively breeding meerkat, female helpers, the philopatric sex, are more responsive to begging than male helpers (25). Understanding which social and physiological factors influence sex differences in parent-offspring communication in avian and non-avian taxa would be illuminating for our understanding of the general (or particular) drivers and outcomes of evolutionary conflicts within the family.

Our study demonstrates the need for comparative studies of the neural and physiological basis of pair bond formation, and how this process intersects with signals from offspring. While numerous studies across species have provided important insights into the neuromolecular underpinnings of pair bond formation and maintenance (e.g. 55–58), we know very little about the mechanisms underlying variation in pair bond strength and mate fidelity or how offspring signals might affect these mechanisms. Similarly, the neural substrates of parental care have been studied in some detail (e.g., 59–61), though how offspring signals modulate these processes is poorly understood (but see 62). However, these studies recommend neuromodulators such as dopamine and the nonapeptides arginine vasopressin and oxytocin as promising candidates, along with the hormone prolactin, likely acting through the medial preoptic area (a central node in social decision-making network across vertebrates 63). Furthermore, glucocorticoid stress hormones can affect pair bond formation and strength, as well as parental care (rodents: 64,65; birds: 66,67). It is thus an intriguing idea that variation in stress physiology may underlie variation in parental decision-making across species. In fact, using non-phylogenetic models, we found that baseline corticosterone levels might mediate how responsive parents are, with a trend for males to become less responsive the higher their baseline stress levels are and for females to be most responsive to offspring at intermediate levels. These results should obviously be considered as very tentative until data from additional species become available that would allow us to confirm whether or not this is a true effect. Overall, our study emphasizes the urgent need for future research in more species on the neural and endocrine mechanisms underlying sexual and parent-offspring conflicts and communication systems.

Surprisingly, we did not find an effect of extra-pair paternity rate, divorce rate, plumage dimorphism, tarsus dimorphism, ecology, or baseline corticosterone levels during nestling care. While it is possible that these factors simply had no impact on parental responsiveness, many of these null results could also be due to a lack of statistical power, since not all data were available for all species. For example, although all 30 species’ pair bond duration could be categorized as typically long-term or short-term, we only had exact divorce rate estimates for 16 species. Life history and hormonal data is invaluable for comparative studies, and many broad evolutionary patterns cannot be understood without such information. As extinction rates soar, obtaining such data from a wide range of species is imperative.

We characterized species as typically having strong or weak pair bonds, according to the classifications in (37). Although mate fidelity and pair bond stability went into this classification, we did not find an effect of divorce rate or extra-pair paternity rates, at either the nestling or brood level. It may be that pair bond type is confounded by some other aspect of a species’ social or abiotic environment. Alternatively, pair bond characteristics may affect responsiveness to begging in a nonlinear fashion, which is not unknown in social evolution (68). Examples for such threshold effects include the lifetime monogamy window in the evolution of eusociality (69); threshold public goods where benefits are only seen after some minimum number of cooperators participate (70); or nonlinear selection pressure of mate choice on male sexual signaling (71). Future work in signaling theory could help disentangle which, if any, aspects of pair bond strength exert the most selective pressure, by varying these independently, and could assess whether these social factors are more likely to have linear or nonlinear impacts on how adults respond to begging signals. Additional empirical data on promiscuity and divorce rates across species would also help clarify these patterns.

In conclusion, our study shows that interactions with offspring mediate maternal and paternal effort, via sex differences in which parent respond more to offspring signals. The direction of this sex difference is associated with a species’ typical pair bond strength: both males and females change their responsiveness to begging when the stability and/or fidelity of bonds differ. Communication between a parent and an offspring does not evolve in a vacuum, it is influenced by the larger social context in which parents have evolved, and the nuances of this communication system vary by sex. Variation in this communication system may also be a mechanism through which sex differences in parental care are engendered or extinguished.

## Author contributions

SMC designed the study and conducted the meta-analysis. SMC and KW extracted and calculated the effect sizes from the literature. All authors discussed and edited the manuscript.

## Acknowledgements

The authors thank Shailee Shah, Ornela De Gasperin, Lauren O’Connell, Ross DeAngelis, Caitlin Friesen, Isaac Miller-Crews, Jiawei Han and Emily Lessig for their comments on the manuscript. We thank Ashleigh Griffin, Stu West, and Camilla Hinde for conversations regarding sex differences in parental care and begging.

