## Supplemental Information for "Sex differences in parental response to offspring begging are associated with pair bond strength across birds"

**Supplemental Figures**

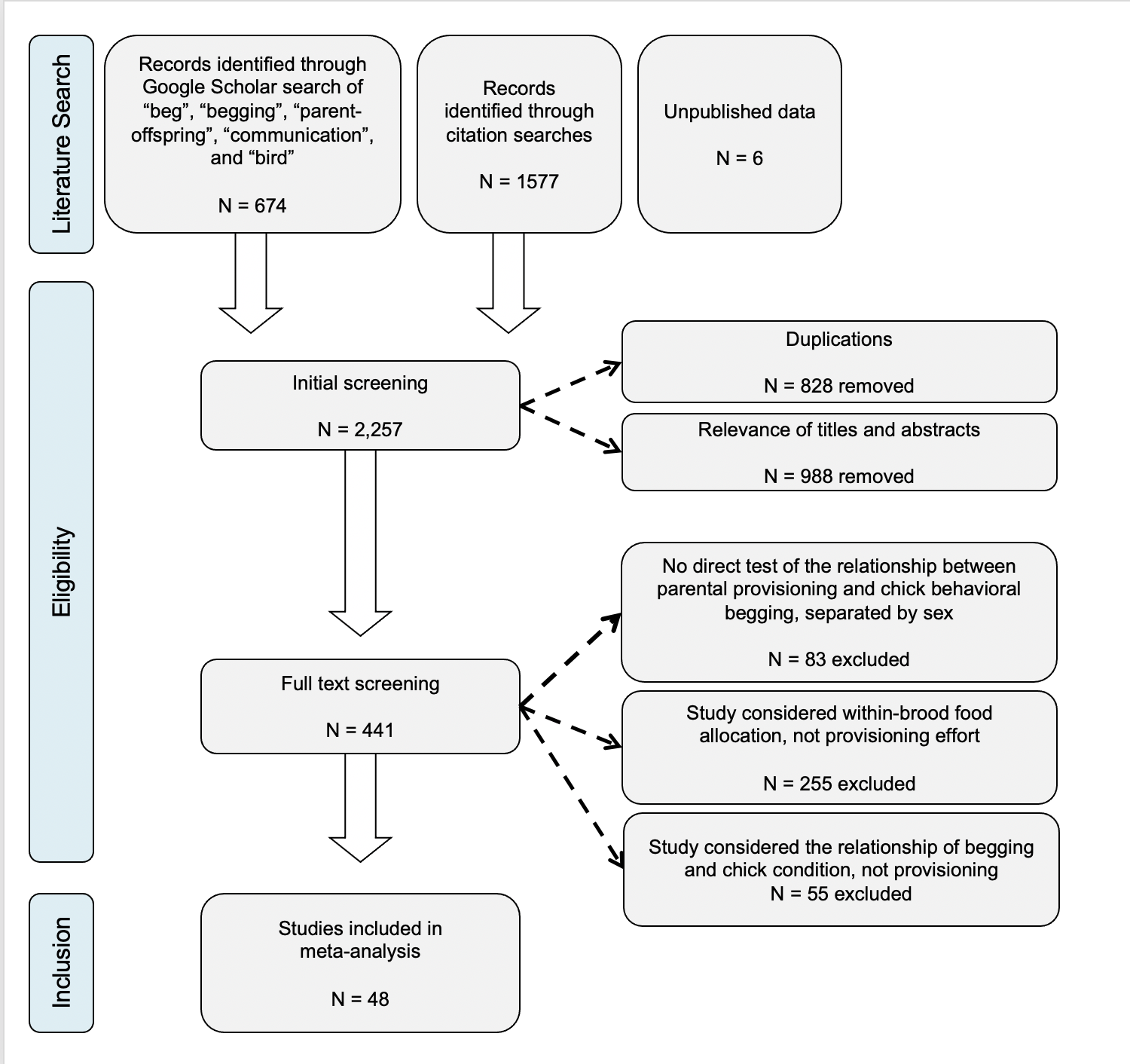

**Supplemental Figure 1.** PRISMA flowchart detailing systematic literature search and exclusion criteria.

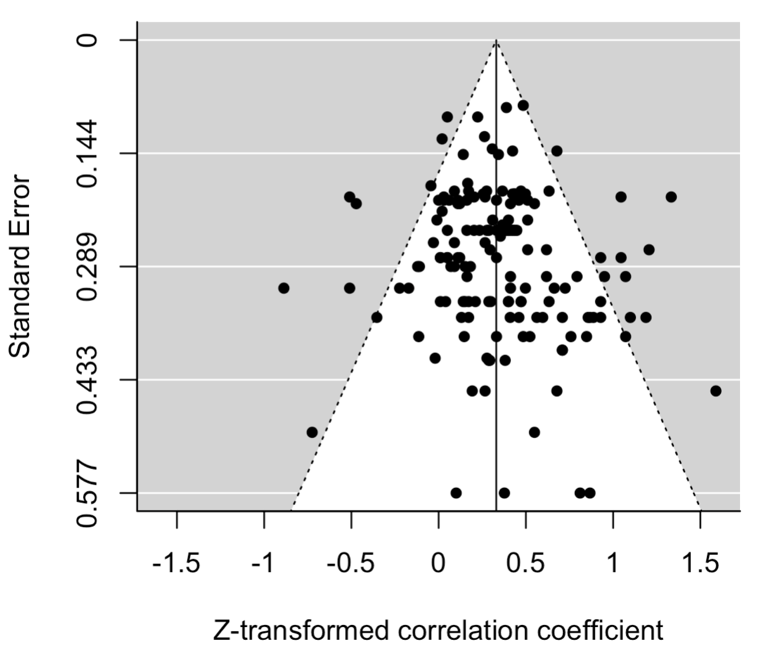

**Supplemental Figure 2.** Funnel plot showing all effect sizes (Z-transformed correlation coefficients) in our across-species dataset (n = 156), compared to their standard error, as calculated based on the sample size in the original test statistic. There is no evidence of publication bias against reporting negative correlations between begging and feeding (Egger’s test: z = 1.28, p = 0.20).

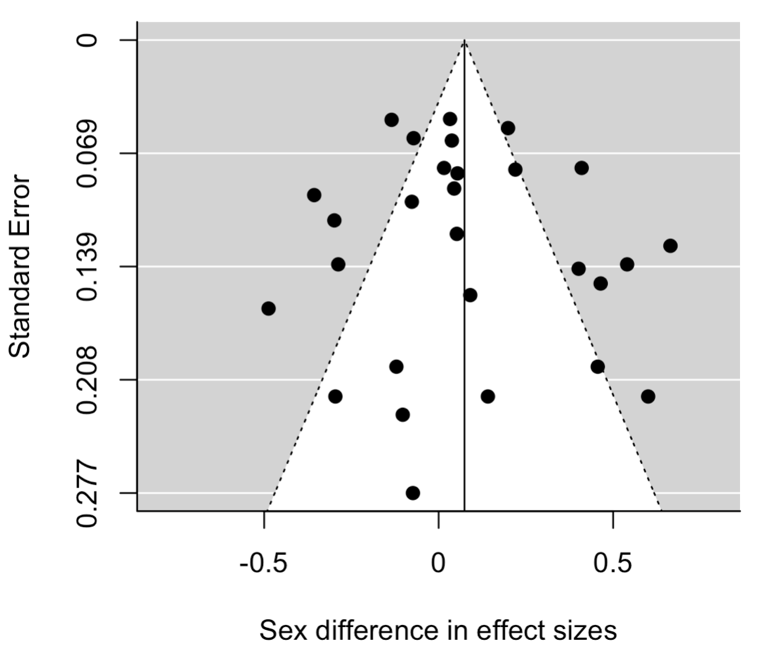

**Supplemental Figure 3.** Funnel plot showing all effect sizes (sex difference) in our within-species dataset (n = 28 species), compared to their standard error, as calculated based on the sample sizes in the original test statistics. There is no evidence of publication bias for sex differences (Egger’s test: z = 0.33, p = 0.74).

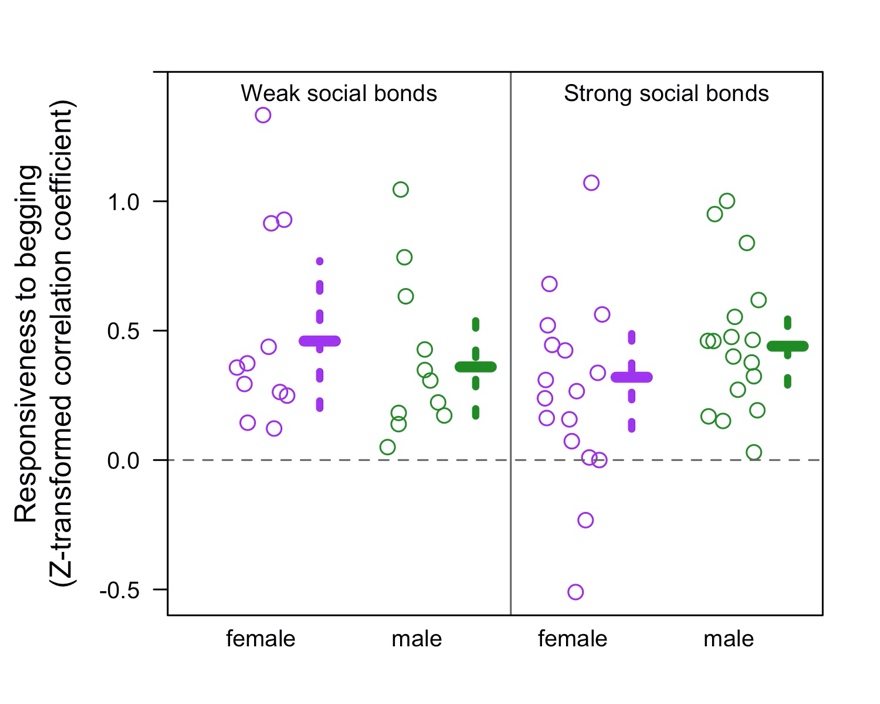

**Supplemental Figure 4.** **Across species, pair bond strength determines how responsive each sex is to begging.** Females are more responsive to begging in species with weak social bonds, while males are more responsive to begging in species with strong social bonds (interaction pMCMC = 0.048*). The y-axis is the Z-transformed correlation coefficient between provisioning and begging. Data points represent each species’ mean effect size for each sex. Solid and dashed lines represent the results from the MCMCglmm models: post.mean and 95% credible intervals. N = 30 species.

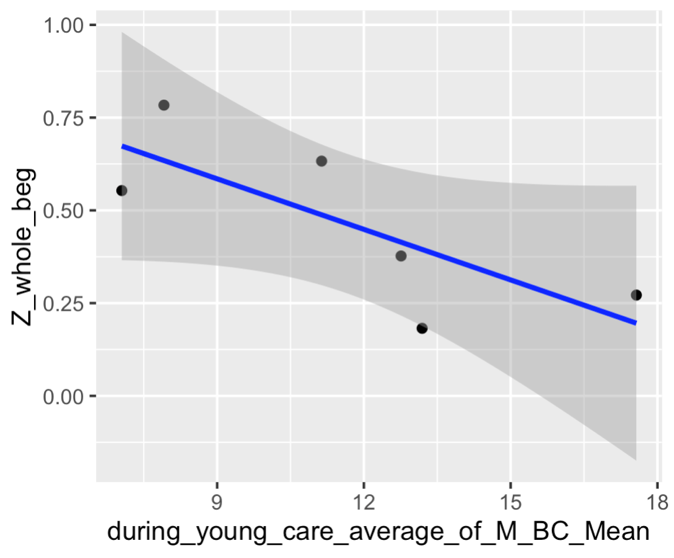

Z-transformed correlation coefficient

Mean baseline corticosterone during young care

(ng/mL, Males)

**Supplemental Figure 5.** Male birds show a trend of decreasing responsiveness to begging in species with higher baseline levels of corticosterone during young care. This effect is not significant when controlling for phylogeny. N = 6 species.

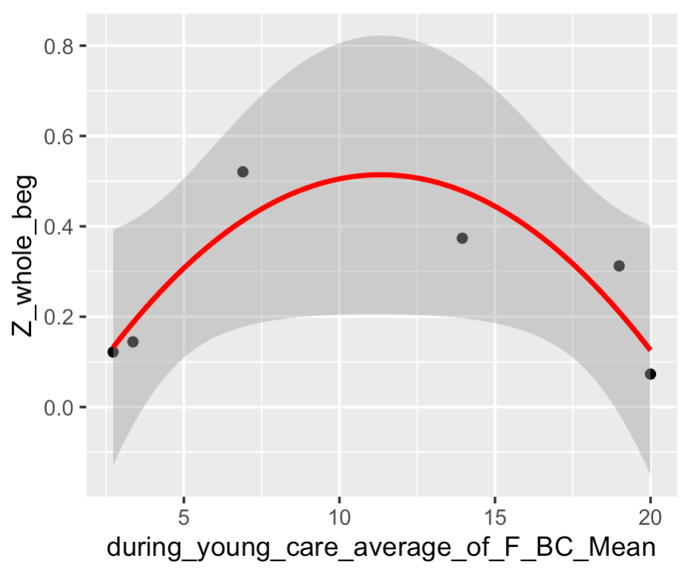

Z-transformed correlation coefficient

Mean baseline corticosterone during young care

(ng/mL, Females)

**Supplemental Figure 6.** Female birds show a trend of being more responsive to begging when the baseline corticosterone levels of the species during young care are at intermediate levels. This effect is not significant when controlling for phylogeny. N = 6 species.

**Supplemental Tables**

|  | **Variable** | **N effect sizes** | **Estimate** | **95% Credible Interval** | **pMCMC** |
| --- | --- | --- | --- | --- | --- |
| **Experiment type** | Experimental | 132 |  |  |  |
|  | Observational | 24 | 0.02 | -0.44 to 0.48 | 0.95 |
| **Estimated** | No | 132 |  |  |  |
|  | Yes | 24 | 0.13 | -0.15 to 0.45 | 0.40 |
| **Beg mode** | Audio | 85 |  |  |  |
|  | Combination | 23 | 0.09 | -0.27 to 0.43 | 0.61 |
|  | Postural | 48 | 0.13 | -0.33 to 0.64 | 0.57 |
| **Beg variable** | Continuous | 113 |  |  |  |
|  | Dichotomous | 43 | -0.03 | -0.27 to 0.25 | 0.86 |
| **Feed variable** | Continuous | 147 |  |  |  |
|  | Dichotomous | 9 | -0.01 | -0.45 to 0.43 | 0.98 |
| **Food deprivation** | Yes | 22 |  |  |  |
|  | No | 134 | 0.04 | -0.18 to 0.30 | 0.70 |
| **Food supplementation** | Yes | 18 |  |  |  |
|  | No | 138 | -0.21 | -0.49 to 0.14 | 0.20 |
| **Brood size manipulation** | Yes | 20 |  |  |  |
|  | No | 136 | 0.18 | -0.27 to 0.65 | 0.45 |
| **Playback** | Yes | 62 |  |  |  |
|  | No | 94 | 0.03 | -0.32 to 0.37 | 0.86 |

**Supplemental Table 3. Potential methodological confounding effects.** Nine models testing for a confounding effect of the original study methodology were run using MCMCglmm with the same conditions and priors as the models in the main text. There is no evidence that study design or the kind of test statistics reported influenced effect size.

|  | **Posterior**  **mean Z** | **95%**  **Credible Interval** | **pMCMC** |
| --- | --- | --- | --- |
| **Fixed effects** |  |  |  |
| (Intercept) | 0.50 | -0.14 to 1.14 | 0.12 |
| Social bond | -0.02 | -0.57 to 0.52 | 0.94 |
| Sex | 0.08 | -0.03 to 0.20 | 0.16 |
| **Social bond * Sex** | **-0.16** | **-0.32 to -0.002** | **0.048*** |
| **Random effects** |  |  | **Variance explained** |
| Phylogeny | 0.25 | 0.09 to 0.46 | 16% |
| Study | 0.14 | 0.07 to 0.23 | 9% |
| Species | 0.20 | 0.08 to 0.34 | 12% |
| Units | 0.004 | 0.00 to 0.01 | 0.2% |

**Supplemental Table 4. How responsive males and females are to begging depends on the type of pair bonds in that species.** Mean results of MCMCglmm analyses on Fisher’s Z-transformed correlation coefficient between behavioral begging and total parental provisioning. Models were generated on 20 random phylogenies, controlling for phylogeny, study and species, and were weighted by the sample size from the original study. Sample error variance was therefore constrained to 1. N = 30 species, 48 studies and 156 effect sizes.

|  | **Posterior**  **mean Z** | **95%**  **Credible Interval** | **pMCMC** |
| --- | --- | --- | --- |
| **Fixed effects** |  |  |  |
| (Intercept) | -0.11 | -0.53 to 0.31 | 0.61 |
| **Social bond** | **0.26** | **0.004 to 0.53** | **0.047*** |
| **Random effects** |  |  | **Variance explained** |
| Phylogeny | 0.21 | 0.09 to 0.35 | 17% |
| Units | 0.01 | 0.00 to 0.04 | 0.1% |

**Supplemental Table 5. Whether males or females of the same species respond more to begging depends on the type of pair bonds in that species.** Mean results of MCMCglmm analyses on the within-species sex difference in Fisher’s Z-transformed correlation coefficient between behavioral begging and total parental provisioning. Negative values indicate females respond more, and positive values indicate males respond more. Models were generated on 20 random phylogenies, and were weighted by the sample sizes for that sex per species in the original studies. Sample error variance was therefore constrained to 1. N = 28 species, data originally from 46 studies and 153 effect sizes.

|  |  |  | **Controlling for phylogeny** | | **Not controlling for phylogeny** | |
| --- | --- | --- | --- | --- | --- | --- |
| **Model** | **N Species** | **Effect** | **Post.Mean** | **pMCMC** | **Estimate** | **p-value** |
| 0 | 30 spp | (Intercept only) | **0.47** | **0.034*** | **0.38** | **<0.0001*** |
| 1 | 30 spp | Sex | 0.02 | 0.71 | *-0.09* | *0.09.* |
|  |  | Males | **0.46** | **< 0.0001** |  |  |
|  |  | Females | **0.43** | **< 0.0001** |  |  |
| 2 | 30 spp | Sex | 0.08 | 0.16 | 0.02 | 0.82 |
|  |  | Social Bond | -0.02 | 0.94 | 0.07 | 0.51 |
|  |  | **Sex * Social Bond** | **-0.16** | **0.048*** | *-0.20* | *0.057.* |
| 3 | 30 spp | Reduction | 0.02 | 0.92 | 0.06 | 0.56 |
|  |  | Environment | -0.04 | 0.65 | -0.07 | 0.46 |
|  |  | Sex | 0.04 | 0.47 | -0.11 | 0.11 |
|  |  | Sex * reduction | 0.02 | 0.77 | -0.03 | 0.81 |
|  |  | Sex * environment | -0.08 | 0.45 | -0.09 | 0.421 |
| 4 | 30 spp | Reduction | 0.04 | 0.87 | 0.06 | 0.72 |
|  |  | Environment | -0.08 | 0.16 | **-0.13** | **0.038*** |
| 5 | 30 spp | Sex | 0.05 | 0.52 | 0.00 | 0.98 |
|  |  | Social Bond | -0.01 | 0.96 | 0.09 | 0.44 |
|  |  | Reduction | 0.04 | 0.88 | 0.05 | 0.68 |
|  |  | Environment | -0.05 | 0.59 | -0.07 | 0.41 |
|  |  | *Sex * Social Bond* | *-0.16* | *0.057.* | **-0.24** | **0.027*** |
|  |  | Sex * Reduction | 0.01 | 0.93 | 0.00 | 0.99 |
|  |  | Sex * Environment | -0.07 | 0.52 | -0.09 | 0.42 |
| 6 | 24 spp | Sex | -0.36 | 0.75 | 1.49 | 0.2 |
|  |  | Tarsus dimorphism | 1.95 | 0.52 | **2.73** | **0.01*** |
|  |  | Sex * tarsus dimorphism | -0.37 | 0.73 | -1.57 | 0.16 |
| 7 | 21 spp | Sex | -0.02 | 0.73 | -0.14 | 0.12 |
|  |  | EPP | 0.00 | 0.99 | 0.00 | 0.54 |
|  |  | Sex * EPP | -0.00 | 0.90 | 0.00 | 0.86 |
| 8 | 16 spp | Sex | -0.09 | 0.45 | 0.10 | 0.50 |
|  |  | Divorce rate | 0.31 | 0.70 | 0.43 | 0.09 |
|  |  | Sex * Divorce rate | -0.21 | 0.36 | -0.44 | 0.11 |
| 9 | 13 spp | Sex | 0.00 | 0.97 | -0.07 | 0.28 |
|  |  | Plumage dimorphism | 0.01 | 0.75 | 0.11 | 0.28 |
|  |  | Sex * Plumage dimorphism | -0.01 | 0.84 | *-0.11* | *0.098.* |
| 10 | 6 spp | Males: Baseline corticosterone levels during young care | -0.03 | 0.8 | *-2.4* | *0.078* |
| 11 | 6 spp | Females: Baseline corticosterone levels during young care | 0.12 | 0.6 | *CORT: 2.97*  *CORT^2^=-3.0* | *0.059*  *0.057* |

**Supplemental Table 6.** Model summaries for all models run. Phylogenetic models were run using MCMCglmm, and controlled for phylogeny, study and species, and were weighted by sample size. Models were run on one random tree. Non-phylogenetic models were run using lme4, and controlled for species. Bold indicates significance at the p < 0.05. Italics indicate significance at the p < 0.10 level.

**Example R code:**

any_sex_difference_erikson_1 <- MCMCglmm(Z_whole_beg ~ which_parent,

random = ~ animal + study + common_name,

prior = list(R=list(V = 1, nu = 0.002), G=list(G1 = list(V=1,nu=2), G2 = list(V=1,nu=2), G3=list(V=1,nu=2))),

pedigree = erikson_30_trees[[1]], mev = full_data$variance ,

data = full_data, family = "gaussian",

verbose=FALSE, pr=TRUE, slice=TRUE,

nitt=3000000, burnin=1000000, thin=1000)

across_species_model_erikson_1 <- MCMCglmm(Z_whole_beg ~ which_parent * social.bondTobias2016,

random = ~ animal + study + common_name,

prior = list(R=list(V = 1, nu = 0.002), G=list(G1 = list(V=1,nu=2), G2 = list(V=1,nu=2), G3=list(V=1,nu=2))),

pedigree = erikson_30_trees[[1]], mev = full_data$variance ,

data = full_data, family = "gaussian",

verbose=FALSE, pr=TRUE, slice=TRUE,

nitt=3000000, burnin=1000000, thin=1000)

within_species_model_erikson_1 <- MCMCglmm(sex_difference ~ social.bondTobias2016,

random = ~ animal,

prior = list(R=list(V = 1, nu = 0.002), G=list(G1 = list(V=1,nu=2))),

pedigree = erikson_30_trees[[1]], mev = data_species_level$variance ,

data = data_species_level, family = "gaussian",

verbose=FALSE, pr=TRUE, slice=TRUE,

nitt=3000000, burnin=1000000, thin=1000)

confounding_factor_playback <- MCMCglmm(Z_whole_beg ~ playback,

random = ~ animal + study + common_name,

prior = list(R=list(V = 1, nu = 0.002), G=list(G1 = list(V=1,nu=2), G2 = list(V=1,nu=2), G3=list(V=1,nu=2))),

pedigree = erikson_30_trees[[1]], mev = full_data$variance ,

data = full_data, family = "gaussian",

verbose=FALSE, pr=TRUE, slice=TRUE,

nitt=3000000, burnin=1000000, thin=1000)
